# Reconfiguration of Functional Brain Hierarchy in Schizophrenia

**DOI:** 10.1101/2024.08.27.608945

**Authors:** Irene Acero-Pousa, Anira Escrichs, Paulina Clara Dagnino, Yonatan Sanz Perl, Morten L. Kringelbach, Peter J. Uhlhass, Gustavo Deco

## Abstract

The multidimensional nature of schizophrenia requires a comprehensive exploration of the functional and structural brain networks. While prior research has provided valuable insights into these aspects, our study goes a step further to investigate the reconfiguration of the hierarchy of brain dynamics, which can help understand how brain regions interact and coordinate in schizophrenia. We applied an innovative thermodynamic framework, which allows for a quantification of the degree of functional hierarchical organization by analysing resting state fMRI-data. Our findings reveal increased hierarchical organization at the whole-brain level and within specific resting-state networks in individuals with schizophrenia, which correlated with negative symptoms, positive formal thought disorder and apathy. Moreover, using a machine learning approach, we showed that hierarchy measures allow a robust diagnostic separation between healthy controls and schizophrenia patients. Thus, our findings provide new insights into the nature of functional connectivity anomalies in schizophrenia, suggesting that they could be caused by the breakdown of the functional orchestration of brain dynamics.

## Introduction

Schizophrenia is a severe and often chronic psychiatric disorder characterized by a range of multidimensional symptoms, including positive symptoms, such as hallucinations and delusions, and negative symptoms, such as anhedonia and avolition (1). The multifaceted nature of schizophrenia is further compounded by fluctuating symptomatology throughout individuals’ lives, leading to significant heterogeneity among patients (2). This complexity and variability require a comprehensive exploration of both the anatomical and functional brain networks.

The presence of neuroanatomical anomalies in both grey and white matter is a well-established phenomenon in schizophrenia (3). Robust evidence has revealed regionally decreased grey matter in both first-episode and chronic schizophrenia patients, specifically in the frontal lobe (4-6), cingulate cortex (4, 7), the thalamus (8, 9), insula (10), post-central gyrus (4), and medial temporal regions (4, 5). In terms of white matter alterations, reductions in the corpus callosum (7), as well as in the frontal and temporal lobes (7, 11) have been extensively documented.

Furthermore, alterations in functional brain dynamics in schizophrenia have been thoroughly reported using resting-state functional magnetic resonance imaging (fMRI) (12, 13). Among the networks considered most relevant are the salience network and the default mode network (DMN) (14, 15). Studies have identified heterogeneous findings, with some indicating hyperconnectivity (16, 17), while others report hypoconnectivity (18, 19) which can vary across networks (19-21).

Despite the numerous identified structural and functional abnormalities, the underlying nature of the pathophysiology remains unclear (22), emphasizing the need for alternative perspectives and hypotheses. One promising approach to conceptualize these anomalies is to consider these deficits as a disorder of integration of information across brain regions (23-26). Information integration can be studied by analyzing the level of asymmetry in the interactions among brain regions, that is, the extent to which information is transmitted in a non-reciprocal manner between them. In systems characterized by symmetrical communication, all regions maintain equal information processing, devoid of inherent hierarchy, thus maintaining a state of equilibrium. On the other hand, asymmetrical interactions indicate distinct information processing, suggesting a hierarchical organization among regions, where certain regions exert greater influence over others (27). This hierarchical arrangement is indicative of non-equilibrium systems, in which some regions assume pivotal roles, orchestrating and exerting a substantial impact on the overall functional brain dynamics (28). Such organization facilitates distinct processing of specialized functional domains and allows for dynamic configuration and intercommunication among brain regions (29, 30).

Hierarchical organization in fMRI-data has been previously assessed in several studies. For instance, previous studies have revealed that states of reduced consciousness exhibit lower hierarchical arrangements compared to conscious wakefulness (31, 32). Moreover, physically and cognitively demanding tasks increase the hierarchical organization compared to a resting state (33, 34). In the context of schizophrenia, some studies suggest overall altered hierarchy (35-39), while others highlight specific alterations in individual networks (40-42).

Here, we apply a new theoretical framework able to quantify the hierarchical organization of brain networks in schizophrenia (43). Specifically, this approach assesses the level of hierarchical organization by quantifying the degree of non-equilibrium in a system, using the fluctuation-dissipation theorem (FDT) and whole-brain modelling (44, 45). Additionally, we explore the correlation between hierarchical organization and symptomatology in schizophrenia. We hypothesize that in schizophrenia, the information flow mediating communication between different regions across the whole-brain is distorted, resulting thus in a malfunctional orchestration of the brain dynamics sustaining normal brain functions. In other words, schizophrenia results from a change of the functional hierarchical brain organization. Moreover, we hypothesize that the symptomatology presented in each individual may correlate with distinct level of hierarchical organization.

## Results

Our study applies a new model-based thermodynamic framework that builds upon the FDT, offering a unique perspective on non-equilibrium brain dynamics and, ultimately, to the hierarchical organization of the brain (43) (**Figure 1c**). The FDT establishes a relationship between the response of a perturbed system to an external force and the internal fluctuations of the same system in the absence of the disturbance (45). A derivative of this theorem, proposed by Onsager, explains that when a system transitions from an initial to a final equilibrium state under a weak external force, this shift can be expressed as spontaneous equilibrium fluctuations. Consequently, these spontaneous fluctuations forecast the dissipation following the perturbation. Nevertheless, in a non-equilibrium system, these intrinsic fluctuations are inaccurate in predicting the dissipation post-perturbation, leading to a violation of the FDT (44) (**Figure 1b**). Hence, an elevated deviation from the FDT signifies a greater degree of non-equilibrium, reflecting more asymmetric interactions between states and, consequently, a heightened level of hierarchical organization (**Figure 1a**).

**Figure 1:**
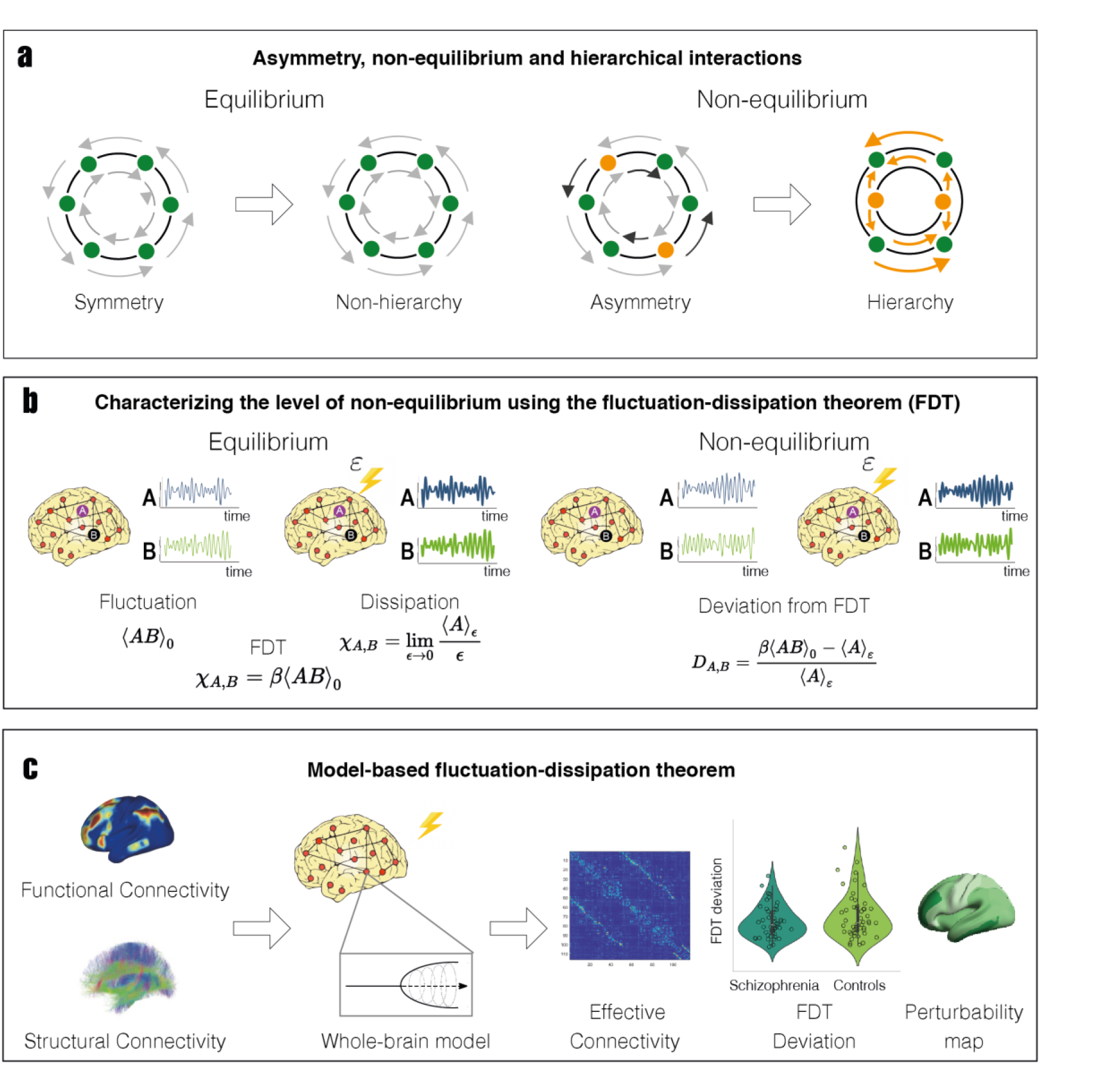
Capturing the hierarchical organization using the FDT. **a)** Hierarchical level is determined by the level of asymmetry of causal interactions between brain regions arising from the breaking of the detailed balance in a system. In equilibrium, the interaction of brain regions is symmetrical, that is, information flows in a reciprocal manner. These symmetrical relationships are in detailed balance, leading to a non-hierarchical organization. In contrast, in non-equilibrium, the asymmetrical interactions break the detailed balance, introducing hierarchical organization in the system. **b)** The level of non-equilibrium can be measured by the deviation of FDT, which can then be employed to characterize the hierarchical organization. In an equilibrium system, spontaneous fluctuations predict the dissipation following the perturbation. Nevertheless, in a non-equilibrium system, the intrinsic fluctuations are not able to forecast the dissipation after a perturbation, leading to a violation of the FDT. **c)** FDT is combined with a whole-brain model fitted to empirical neuroimaging data, incorporating functional and structural connectivity. Each node’s local dynamics of the model is described as the normal form of a supercritical Hopf bifurcation. The optimized model provides the effective connectivity, the FDT deviations as well as the perturbability maps for different brain states.

### Hierarchical reconfiguration across the whole-brain

We initially assessed the difference in hierarchical organization between schizophrenia and control groups at the whole-brain level by leveraging the level of FDT deviations in each participant. At the whole brain-level, group differences reached a trend-level (p = 0.058, median (interquartile range (IQR)): schizophrenia = 28.65 (16.02), controls = 27.08 (14.72)), and the distribution of values indicates a higher level of FDT violations in schizophrenia (**Figure 2a**, left panel). FDT violations were significantly increased in schizophrenia patients when only cortical areas were considered (p < 0.05, median (IQR): schizophrenia = 29.53 (16.33), controls = 28.26 (16.25)), (**Figure 2a**, middle panel). In a subsequent analysis, we investigated whether the observed significance in cortical areas could also be attributed to other nodes. For each subject, we averaged the FDT values of the nodes exhibiting the highest differences between the two groups, only excluding the five most similar ones, and found a statistically significant increase in FDT violations in the schizophrenia group (p < 0.05, median (IQR): schizophrenia = 29.02 (16.19), controls = 27.04 (14.61), (**Figure 2a**, right panel), suggesting an overall increase in hierarchical brain organization in comparison to the control group.

**Figure 2:**
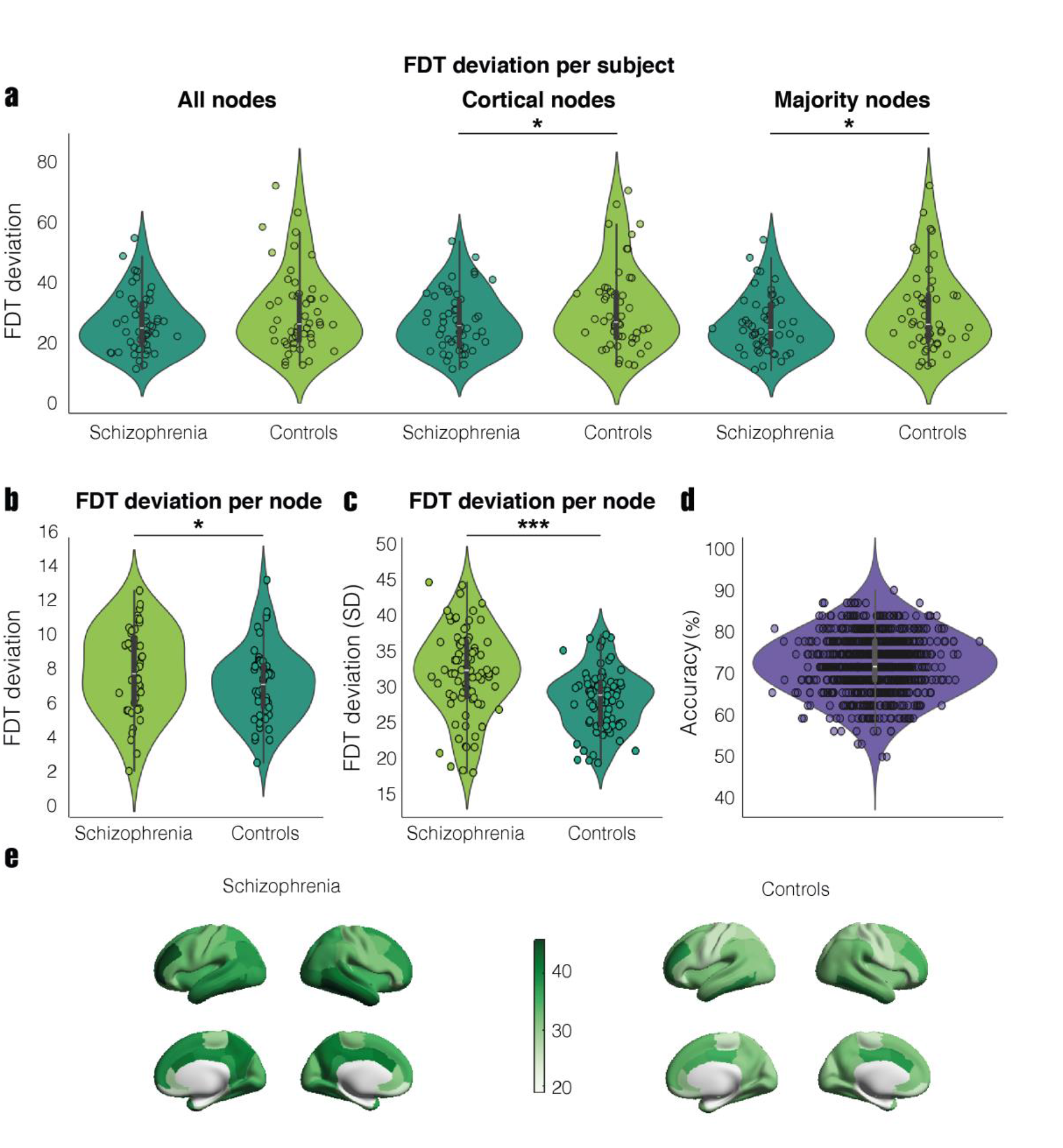
FDT deviation values across the whole-brain. Analysis of the whole-brain FDT deviation values between schizophrenia and control patients, to understand differences in hierarchical brain organization. **a)** Boxplots illustrate the distribution of the average FDT values per subject. The “All nodes” panel presents the analysis of all nodes, the “Cortical nodes” panel exclusively includes the cortical nodes and the “Majority nodes” panel includes nodes with the highest differences between groups, excluding the five most similar. The findings reveal statistically significant elevated values in schizophrenia compared to the control group in the “Cortical nodes” and “Majority nodes” panels (p < 0.05), while results in the “All nodes” panel are not significant (p = 0.058). **b)** Boxplots show the distribution of the average FDT values per node. Results indicate higher statistically significant values in schizophrenia compared to controls (p < 0.001). **c)** Boxplots depict the distribution of the standard deviation of the average FDT values across nodes. Results indicate higher statistically significant variation in schizophrenia compared to controls (p < 0.001). **d)** The violin plot displays the distribution of the support vector machine model’s accuracy across 1000 k-folds using the FDT violations of each subject as input. **e)** Brain renders represent the spatial distribution of the average FDT values per node for each group. Asterisks in the figure indicate statistical significance: * represents p < 0.05 and *** represents p < 0.001.

Additionally, we explored the hierarchical organization for each brain area instead of each subject, aiming to provide insights into the specific regions implicated in the observed differences and mitigate the potential confounding effects of individual variability. We computed the degree of FDT deviation for each brain area (i.e., for each group, the average value across all subjects for each brain region) (**see Figure 2e**). Statistical comparisons demonstrated a significantly increased level of FDT violations in schizophrenia (p < 0.001, median (IQR): schizophrenia = 33.36 (7.85), controls = 29.77 (5.76)) (**Figure 2b**). Subsequently, we examined the consistency of FDT deviation values among brain areas within each group to determine whether the elevated FDT deviation was associated with specific regions or reflected a more generalized pattern. For this analysis, we calculated the standard deviation (SD) of FDT deviation values across all regions for each group, revealing higher statistically significant values in schizophrenia (p < 0.001, median (IQR): schizophrenia = 16.66 (2.62), controls = 12.44 (2.64)) (**Figure 2c**). This observation indicates that controls exhibit greater uniformity in the deviation of FDT across brain areas, suggesting that the increased whole-brain FDT values observed in schizophrenia may stem from specific regions with notably higher FDT levels. This localized elevation contributes to the increased variance observed within the schizophrenia group.

To further examine whether FDT violations distinguish between schizophrenia subjects and controls, we developed a support vector machine (SVM) classification model. The model was trained using the average FDT deviation of each subject as input features and achieved an accuracy of 74.46% (**Figure 2d**), indicating that the hierarchical organization can serve as a promising model-based biomarker.

### Hierarchical reconfiguration across resting-state networks

To investigate the localization of the alterations, we investigated the FDT deviations across eight RSNs: the visual, somatomotor, dorsal attention, salience, limbic, control, DMN and subcortical networks. As in the whole-brain assessment, we conducted the analysis both at the subject/single-participant perspective and the brain region perspective.

At the single-participant level, findings reveal a persistent increased FDT deviation in schizophrenia across all RSNs (**Figure 3a**). Statistically significant differences emerge in the visual (p < 0.05), somatomotor (p < 0.05), dorsal attention (p < 0.01) and DMN (p < 0.05). However, no significance is found in the salience (p > 0.05), limbic (p > 0.05), control (p > 0.05) and subcortical (p > 0.05) networks. Only the dorsal attention network remains significant after correcting for multiple comparisons (**see Table 1**). At the brain area level, results also indicate a consistent pattern of elevated FDT deviation within schizophrenia across all RSNs (**Figure 3b**). Notably, statistically significant differences appear in the visual (p < 0.01), somatomotor (p < 0.01), salience (p < 0.05) and DMN (p < 0.01). Results do not indicate significance in the dorsal attention (p > 0.05), limbic (p > 0.05), control (p > 0.05) and subcortical (p > 0.05) networks. All significant p-values persist following correction for multiple comparisons (**see Table 2**).

**Table 1:** Statistical metrics of the FDT deviation values across RSNs from the subject perspective. This table presents the median and interquartile range (IQR) of the average FDT deviation values per subject in each group, categorized by RSNs. The significance column indicates the p-values obtained, with black asterisks indicating statistical significance after FDR correction, red asterisks representing the significance prior to FDR correction, indicating they do not survive the correction.

| RSN | FDT deviations in schizophrenia: median (IQR) | FDT deviations in controls: median (IQR) | Significance (p) |
| --- | --- | --- | --- |
| Visual | 31.01 (18.61) | 27.44 (16.87) | $p < 0.05$ * |
| Somatomotor | 27.05 (21.05) | 24.88 (10.63) | $p < 0.05$ * |
| Dorsal Attention | 30.78 (18.86) | 26.25 (12.34) | $p < 0.01$ ** |
| Salience | 33.03 (18.87) | 32.28 (17.17) | $p > 0.05$ |
| Limbic | 29.89 (14.52) | 25.34 (16.41) | $p > 0.05$ |
| Control | 32.79 (18.92) | 31.72 (15.94) | $p > 0.05$ |
| DMN | 29.37 (19.25) | 28.24 (17.04) | $p < 0.05$ * |
| Subcortical | 24.95 (15.66) | 23.43 (10.53) | $p > 0.05$ |

**Table 2:**
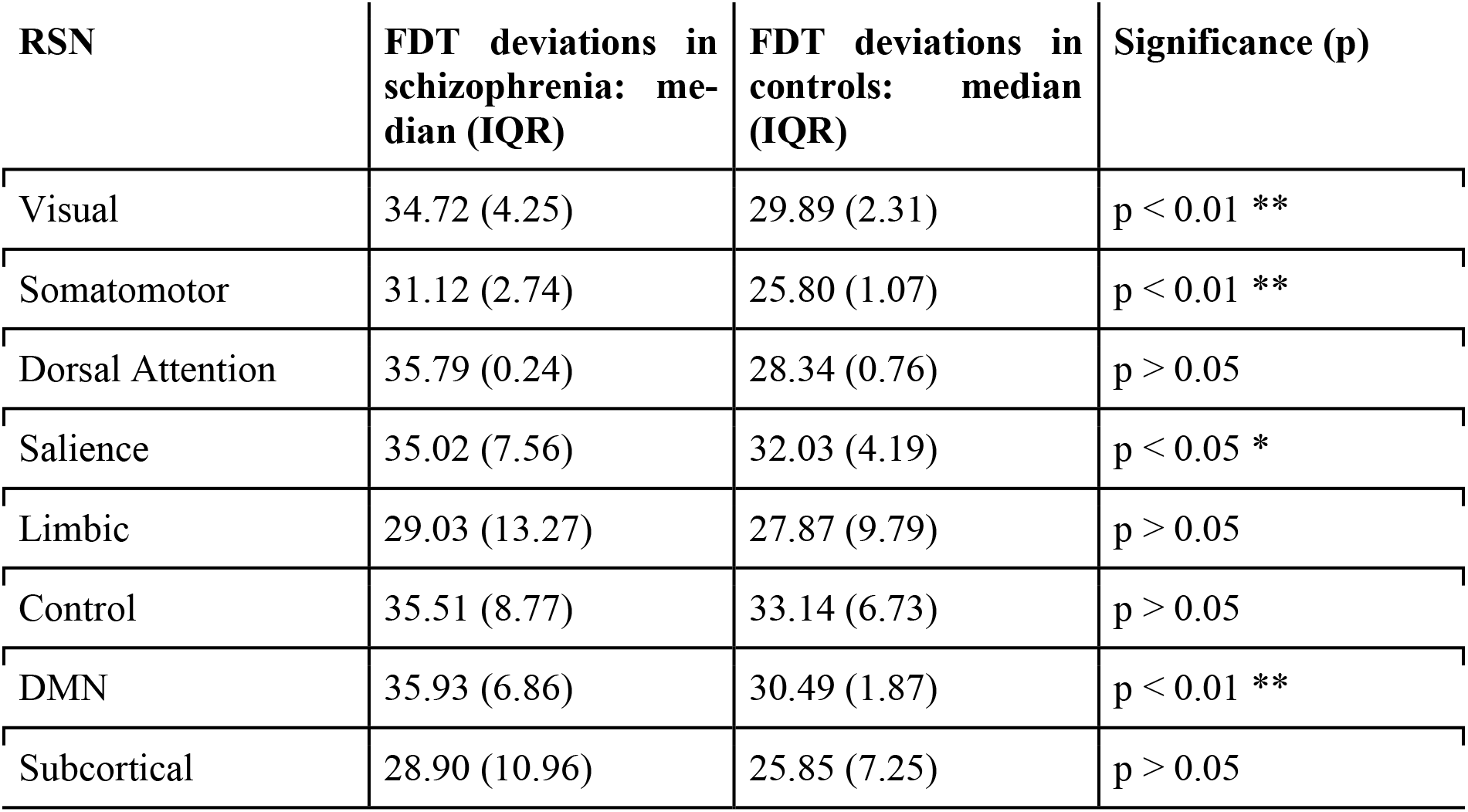
Statistical metrics of the FDT deviation values across RSN from the node perspective. This table indicates the median and interquartile range (IQR) of the average FDT deviation values per node in each group, categorized by RSNs. The significance column indicates the p-value obtained after FDR correction.

**Figure 3:**
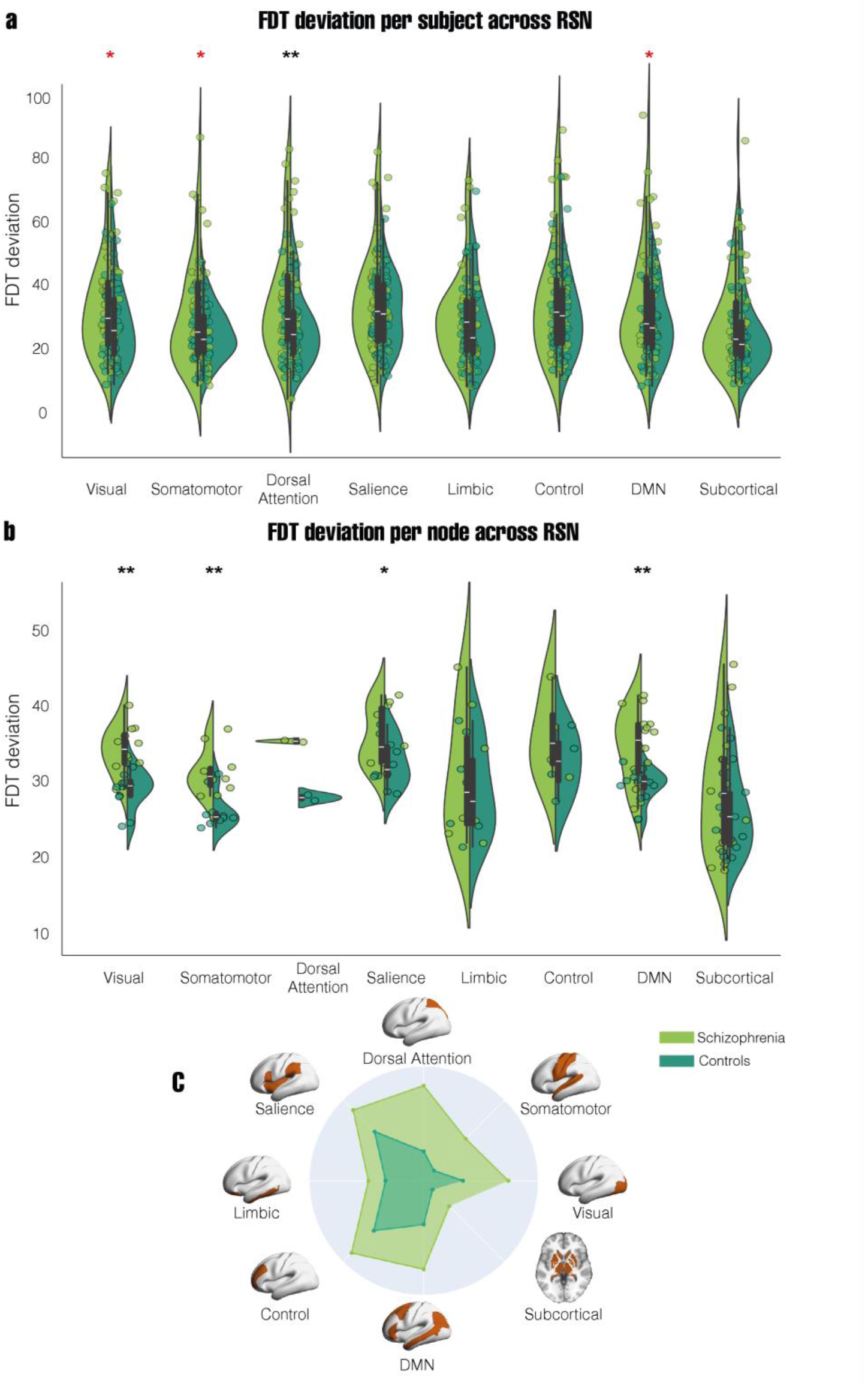
FDT deviation values across RSNs. Comparison of FDT deviation values between schizophrenia and control groups to identify the networks contributing to the difference in hierarchical organization. **a)** Boxplots illustrate the distribution of the average FDT values per subject for each RSN. Results indicate higher median values in schizophrenia in all networks, with significant statistical differences in the visual (p < 0.05), somatomotor (p < 0.05), dorsal attention (p < 0.01) and DMN (p < 0.05). **b)** Boxplots illustrate the distribution of the average FDT values per node for each RSN. Results indicate higher median values in schizophrenia in all networks, with significant statistical differences in the visual (p < 0.01), somatomotor (p < 0.01), salience (p < 0.05) and DMN (p < 0.01). **c)** Radar plot shows the mean FDT values per RSN for each group on the same scale, complemented with a brain render highlighting in orange the area of the corresponding network. Black asterisks in the figure indicate comparisons that remain significant after FDR correction, while red asterisks indicate comparisons that do not. * represents p < 0.05 and ** represents p < 0.01.

We represented the mean FDT deviation values of all subjects for each RSN and each group on a uniform scale, aiming to identify the regions with the largest differences in hierarchical organization between the two groups (**Figure 3c**). The findings indicate that the regions orchestrating the changes in hierarchical configuration are the dorsal attention (standardized mean difference (SMD) = 0.54), somatomotor (SMD = 0.41), visual (SMD = 0.38) and DMN (SMD = 0.36), followed by the control (SMD = 0.26), salience (SMD = 0.23), subcortical (SMD = 0.21) and finally limbic (SMD = 0.15) networks (**Table 3**).

**Table 3:** Statistical metrics of the FDT deviation values across RSN for the comparison between groups. This table indicates the mean and standard deviation (SD) of the average FDT deviation values per subject in each group, by RSNs. It also indicates the difference between the two groups, computed by subtracting their mean values and dividing by the standard deviations.

| RSN | FDT deviations in schizophrenia: mean (SD) | FDT deviations in controls: mean (SD) | Standardized mean difference between groups |
| --- | --- | --- | --- |
| Visual | 34.61 (14.36) | 29.45 (12.02) | 0.38 |
| Somatomotor | 31.74 (15.67) | 26.64 (7.67) | 0.41 |
| Dorsal Attention | 35.79 (16.69) | 28.33 (10.11) | 0.54 |
| Salience | 36.34 (15.30) | 32.88 (10.35) | 0.26 |
| Limbic | 31.28 (13.28) | 29.34 (12.41) | 0.15 |
| Control | 36.59 (17.10) | 33.01 (12.51) | 0.23 |
| DMN | 35.04 (16.32) | 29.96 (10.95) | 0.36 |
| Subcortical | 29.10 (13.71) | 26.42 (10.69) | 0.21 |

### Hierarchical reconfiguration correlates with symptom severity

Next, we explored the potential correlation between the changes of the hierarchical organization in schizophrenia and specific symptoms. We employed exploratory factor analysis to reduce the dimensionality of the symptom’s data from 58 to 3 factors. The analysis revealed a positive significant correlation between FDT deviations and Factor 1 (related to negative symptoms) (r = 0.33, p < 0.05), a negative significant correlation between FDT deviations and Factor 3 (associated with features of positive formal thought disorder and apathy) (r = -0.31, p < 0.05), but no significant correlation between FDT deviations and Factor 2 (characterized by positive symptoms) (r = -0.23, p > 0.05) (**Figure 4b**).

**Figure 4:**
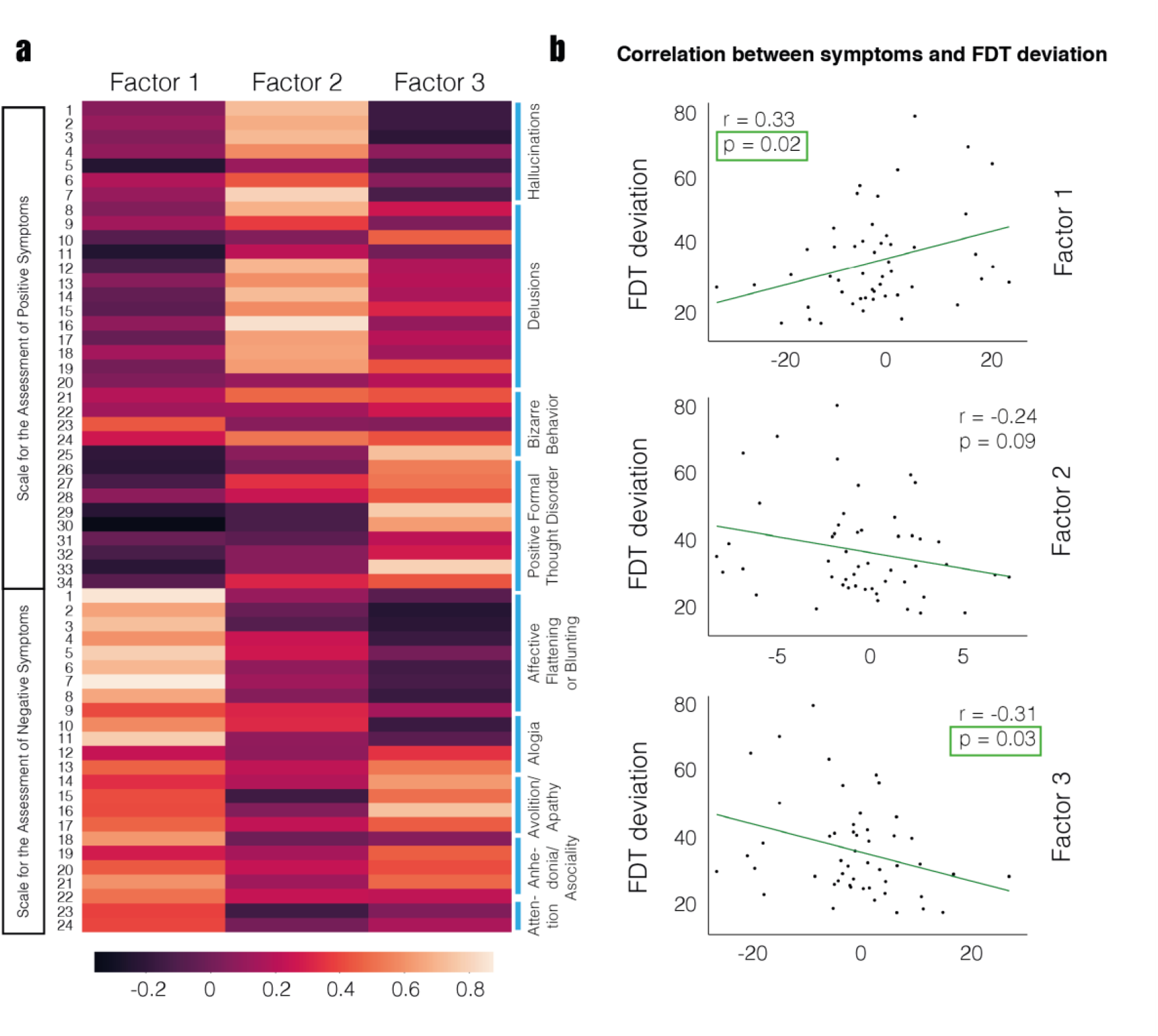
Correlation analysis between FDT deviation and symptom severity. Exploration of the correlation between the FDT deviation value of each subject and their symptomatology items evaluated using the SAPS and SANS scales. **a)** Loading matrix of the three-factor solution derived from the exploratory factor analysis, including SAPS and SANS variables. Each row indicates an item on the scales. Interestingly, Factor 1 shows strong values for the SANS variables, Factor 2 for the SAPS, and Factor 3 for positive formal thought disorder and avolition/apathy items. **b)** Correlation between the factor scores of each subject and their average FDT deviation values. Factor 1 exhibits a significant positive correlation (p < 0.05, r = 0.33) with FDT deviation, while Factor 2 does not demonstrate significant correlation (p > 0.05, r = -0.24) and Factor 3 shows a negative significant correlation (p < 0.05, r = -0.31). This indicates that FDT deviation is correlated with the severity of negative symptoms and items of positive formal thought disorder and avolition/apathy.

### Hierarchical reconfiguration can classify between individuals with schizophrenia and controls

We determined whether differences in brain dynamics between schizophrenia and controls can be more effectively explained due to changes in brain hierarchy (as reflected in FDT deviations) or by other functional metrics in the literature (Functional Connectivity (FC) and Effective Connectivity (EC)). We trained an SVM classifier using the FC, the EC, the FDT deviations and a combination of EC and FDT (EC + FDT) as input. Our results indicate that the EC+FDT input combination achieves the highest accuracy among the classifiers (82.4%), surpassing the 55.38% obtained using FC, 73.12% with EC and 74.02% with FDT alone (**Figure 5a, 5b**).

**Figure 5:**
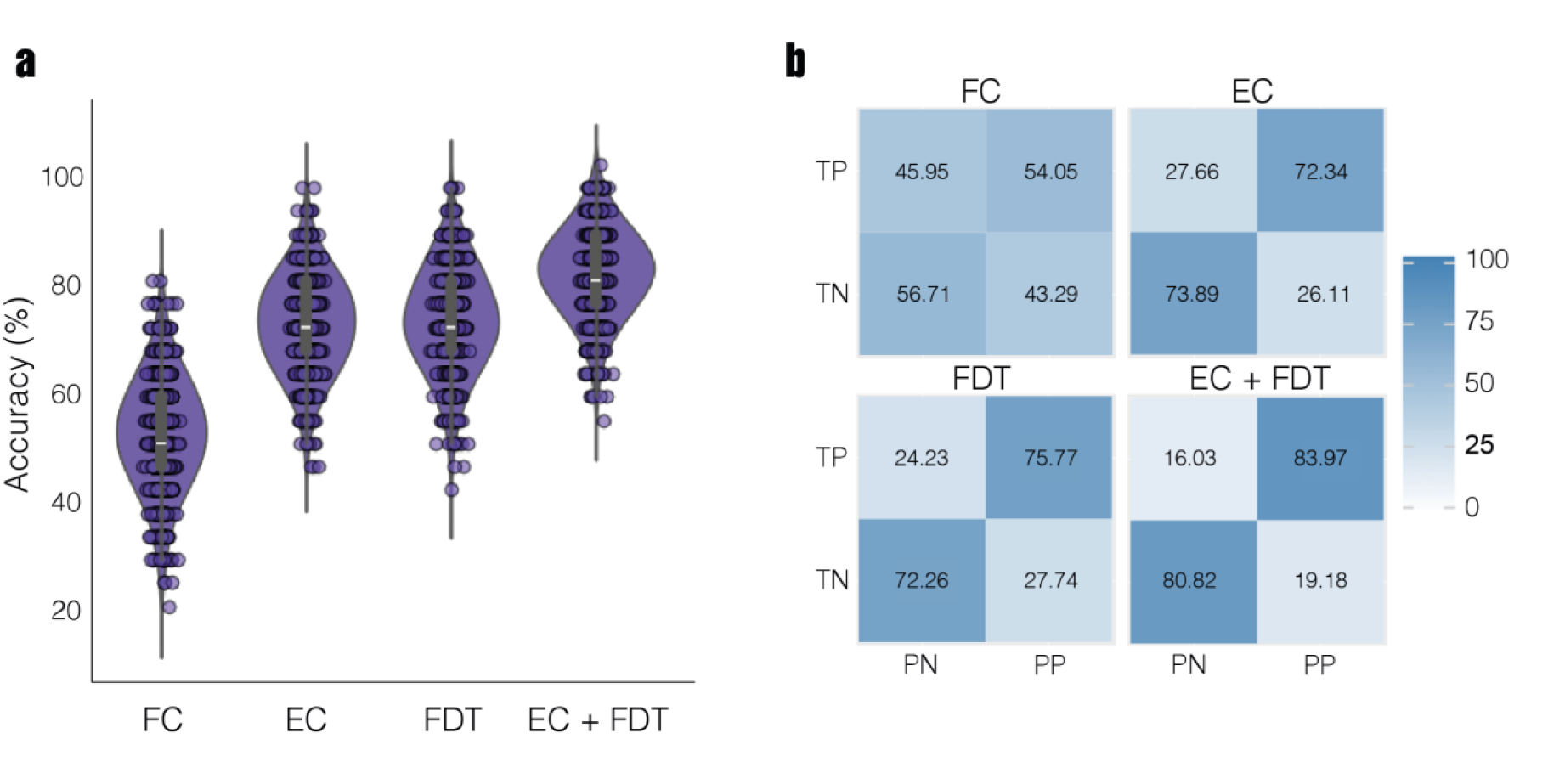
Support Vector Machine classification. Comparison of the accuracy obtained using a SVM algorithm with different inputs (FC, EC, FDT and EC+FDT) to identify the most informative metrics. **a)** Violin plots display the distribution of SVM accuracy across 1000 k-folds using FC, EC, FDT and EC + FDT data as inputs. **b)** Confusion matrices illustrate the overall accuracy achieved corresponding to the distributions in (a), presented as percentages represented on a color scale. TP indicates true positive, TN indicates true negative, PN indicates predicted negative and PP indicates predicted positive. The average accuracy obtained using FC data was 55.38%, 73.12% with EC, 74.02% with FDT and 82.4% with EC + FDT, indicating that the combined EC and FDT metrics are the most informative for distinguishing between groups.

Finally, focusing on the EC+FDT SVM model, we conducted an in-depth analysis by identifying the 10 percent most influential features for the classification, corresponding to the highest coefficients of the model (**Table 4**). Interestingly, some of the nodes overlap with the significant networks found in the analysis across RSNs (**Figure 3a, b**), that is, the DMN, somatomotor and visual networks. Moreover, the subcortical and limbic networks also appear to be highly important for the classification.

**Table 4:**
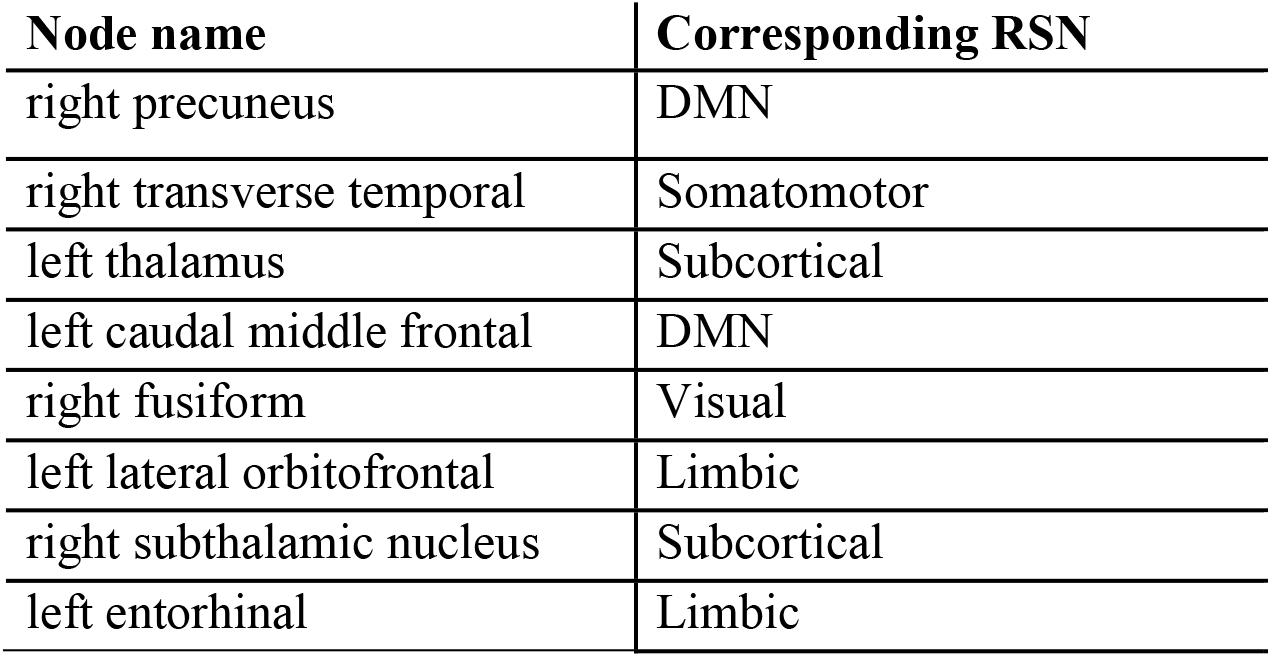
SVM Feature Importance Results. This table presents the 10% most important features obtained through SVM classification, indicating the relevance of specific nodes and their corresponding RSN, sorted in descending order. The higher the feature in the table, the greater the significance of the node in classification.

## Discussion

We integrated a whole-brain computational model with the fluctuation-dissipation theorem (FDT) investigate whether the brain’s hierarchical organization differs in patients with schizophrenia, using FDT deviations as a metric of hierarchy. Our analysis revealed consistently heightened organization in schizophrenia, with significant differences observed in the visual, somatomotor, dorsal attention, DMN and salience networks. Moreover, these changes correlated with the severity of formal thought disorder, negative and positive symptoms. Finally, a SVM model demonstrated that combining FDT and EC classifies with superior performance schizophrenia patients from controls.

At the whole-brain level, we found increased FDT deviation in schizophrenia. A deviation from the FDT is characteristic of a system departing from the equilibrium (44, 45), which is defined by asymmetrical interactions between its components and a hierarchically organized structure (27). Increased FDT deviation suggests an elevated level of non-equilibrium and more hierarchically organized brain dynamics in schizophrenia compared to controls, suggesting a stronger contribution of top-down processing (27). Specifically, we found that the higher levels of the hierarchy are occupied by the visual, somatomotor, salience and DMN networks. These areas exhibit greater influence over the other RSNs, driving the orchestration and organizing the functional dynamics to a higher extent than controls.

Previous studies have also shown differences in the whole-brain hierarchical organization of networks in schizophrenia (37, 38, 46), while others have highlighted reduced hierarchical organization (27, 47, 48). Notably, a previous study found that the connectivity between the DMN with other regions was more consistently present in individuals with schizophrenia than controls (49). Moreover, Yang et al (36) demonstrated that higher functional hierarchy in the DMN underlies the persistently observed elevated FC in studies of schizophrenia (16, 50, 51). Increased connectivity between DMN and both the somatomotor and visual networks was also reported in (52). This goes in line with our outcomes, emphasizing the important role of this network in schizophrenia. In contrast, regarding the other networks (i.e., the visual, somatomotor and salience), previous research points to reduced levels of hierarchy (37, 42). Nevertheless, measures of entropy, which indirectly captures hierarchy (53), has indeed presented increased results in the visual network (21, 54).

Our findings highlight correlations between the hierarchical organization in individuals with schizophrenia and the severity of negative symptoms, positive formal thought disorder and apathy. Previous studies have reported correlations between changes in DMN connectivity and negative symptoms (55, 56). Hierarchical changes within this network could be affecting processes such as self-referential thinking and introspective cognition. Additionally, research has indicated that the somatomotor network can also impact negative symptoms (57). Alterations in its hierarchical organization could impact motor planning and execution, contributing to symptoms such as psychomotor retardation and speech abnormalities. Furthermore, dysregulation of the salience network has been associated with negative symptoms and motivational deficits in schizophrenia (15, 58), potentially leading to apathy and reduced responsiveness to environmental cues. These findings are consistent with our outcomes, suggesting that alterations in the hierarchical organization of specific networks may underlie the manifestation of symptoms commonly observed in the disease.

Finally, we employed a machine learning classification model to evaluate whether the differences in brain dynamics between schizophrenia and controls are better captured by alterations in brain hierarchy or by other functional metrics, such as FC and EC. The model demonstrated the highest performance when trained with a combination of EC and FDT deviations. EC provides information about the causal influence or directed connections between brain regions, indicating how one region influences another. Beyond these, the model also underscored the relevance of the subcortical and limbic networks, which were not prominently identified in the previous statistical analysis. This underscores the complementary nature of machine learning approaches in identifying patterns that may elude traditional statistical methods. This finding is interesting given that disruptions in hierarchy in the subcortical and limbic systems have been reported previously (59, 60).

Our results are potentially compatible with shifts in the balance between excitation (E) and inhibition (I) mediated by NMDA receptor hypofunction implicated in schizophrenia (61-63). Interestingly, previous research in primates using computational models proposed that variations in local recurrent excitation strength at different hierarchical levels account for the observed differences in neural activity time-scales across cortical regions (64-66). To bridge between functional neuroimaging and neuronal-level alterations, a previous study constructed a biophysical model integrating the hierarchical brain organization and E/I perturbations, which was able to predict the brain dynamics observed in the disease (36). Moreover, Brau et al (67) observed the effects of an NMDA antagonist in healthy controls, which led to increased network flexibility (i.e., higher reconfiguration of brain networks). Also using an NMDA antagonist for anaesthesia, Deco et al (32) predicted heightened hierarchy in primates compared to their awake state. As a future step, it would be valuable to extend our framework by including receptor maps, such as in (68, 69), to obtain a deeper and more precise understanding of our findings.

The current study has several limitations. First, although consistent with comparable studies in the field (70, 71), the dataset analysed in this study is relatively small. Second, a potential limitation arises from the conceivable influence of medication on brain dynamics, which could impact the observed brain activity and subsequent results (72). Third, in the parcellation employed in the analysis (i.e., the DK80 atlas (28)), the dorsal attention network is constituted by only two nodes and, thus, we refrain from making detailed interpretations when comparing groups by nodes.

In summary, we used a computational whole-brain model and the violations of the FDT and found elevated hierarchical brain organization in schizophrenia compared to controls both across the wholebrain and RSNs. Furthermore, we found that the changes in hierarchy are correlated with negative symptoms, positive formal thought disorder and apathy. Finally, through SVM-based pattern identification, we demonstrated that combining EC and FDT resulted in superior classification accuracy than individual metrics (i.e., FC, EC and FDT deviations). Overall, this framework can be potentially used as a model-based biomarker to differentiate schizophrenia from healthy controls. Importantly, this inquiry is not limited to schizophrenia alone but opens up broader perspectives for understanding the hierarchical brain organization of other psychiatric conditions and different brain states.

## Methods

### Participants

In this study, we analyzed resting-state fMRI data sourced from the UCLA Consortium for Neuropsychiatric Phenomics LA5c Study ((74), open-neuro.org/datasets/ds000030). Our participant pool encompassed two age-matched cohorts: 50 individuals diagnosed with schizophrenia (12 female, mean group age: 36.5 ± 8.8, standard error of the mean (SEM) ± 1.26) and 50 healthy subjects (16 female, mean group age: 36.6 ± 8.9, SEM ± 0.88). All participants provided written informed consent following the University of California Los Angeles Institutional Review Board’s approved protocols. Exclusion criteria encompassed individuals with multiple diagnoses, substance abuse (excluding caffeine or nicotine), major depressive disorder, suicidality, or anxiety disorders (obsessive-compulsive disorder, panic disorder, generalized anxiety disorder, and post-traumatic stress disorder). Stable medication usage was allowed for patients. Control group inclusion criteria involved individuals with no prior psychiatric diagnoses or familial psychiatric history and no prior exposure to psychotropic drugs. Left-handed participants and those with MRI contraindications were also excluded. Detailed dataset information is available in the data description provided by Poldrack et al (75).

### Functional MRI Data Acquisition

MRI data was acquired using two distinct scanners, the Ahmanson-Lovelace Brain Mapping Center (Siemens version syngo MR B15) and the Staglin Center for Cognitive Neuroscience (Siemens version syngo MR B17) at UCLA. Functional MRI data was collected using a T2* weighted echoplanar imaging sequence with oblique slice orientation (slice thickness = 4 mm, 34 slices, TR = 2 s, TE = 30 ms, flip angle = 90°, matrix 64 × 64, FOV = 192 mm, 152 volumes). An anatomical volume was also captured using a T1-weighted sequence (TR = 1.9 s, TE = 2.26 ms, FOV = 250 mm, matrix = 256 × 256, sagital plane, slice thickness = 1 mm, 176 volumes). Throughout the experiment, participants were instructed to remain relaxed with their eyes open.

Preprocessing of the fMRI data involved fMRIPrep v1.1 (76), a Nipype-based tool developed by Gorgolewski et al (77). To enhance the fMRIPrep outputs, a nonaggressive variant of ICA-AROMA was applied to the unsmoothed data. Additionally, mean white matter and cerebrospinal fluid signal were regressed out and no GSR was utilized during data processing. Finally, the brain was divided using the DK80 parcellation (28), including 62 cortical (31 per hemisphere), and 18 subcortical areas (9 per hemisphere), a total of 80 regions.

### Diffusion MRI Data Acquisition

For structural connectivity data acquisition, we processed multi-shell diffusion-weighted imaging data from 32 participants drawn from the Human Connectome Project (HCP) database. The scans, spanning approximately 89 minutes, adhered to acquisition parameters detailed in Thomas et al (78). Connectivity was estimated using the method outlined by Horn et al (79). Briefly, the data was processed using a generalized q-sampling imaging algorithm implemented in DSI studio (http://dsi-studio.labsolver.org). T2-weighted anatomical images were segmented to generate a white-matter mask, and image co-registration to the b0 image of the diffusion data was performed employing SPM12. Within this refined white-matter mask, we sampled 200,000 fibres for each participant in the Human Connectome Project. These fibres were then transformed into MNI space, utilizing Lead-DBS (80). The methodology was designed to employ algorithms demonstrated to be optimal, minimizing falsepositive fibres, as underscored in recent open challenges (81, 82).

### Theoretical Framework

#### Violations of the Fluctuation-Dissipation Theorem

The Fluctuation-Dissipation Theorem (FDT) elucidates the intricate relationship between a perturbed system’s response to an external force and its internal fluctuations in the absence of such disturbances (45). To quantify the violations of the FDT, we adopt Onsager’s approach, who, through his regression principle, devised a simple derivation of the FDT (44, 83). According to this principle, if a system evolves from an initial equilibrium state 1 to a different equilibrium state 2 due to a weak external force, this transition can be expressed as a spontaneous equilibrium fluctuation. Consider a system in the initial equilibrium state 1, unperturbed, which is at temperature *T* and has a probability distribution of system configuration *G*, named as *P*_0_(*G*). At *t* = 0, an external force *ε* coupled to an observable *B*(*G*) is applied, resulting in a change in the energy of the system and transforming the initial state 1 towards the final state 2. We can understand the impact of the perturbation applied in *B* by observing the expectation value of a second observable ⟨*A*(*t*)⟩_*ε*_ coupled to an observable *A*(*G*) from the initial unperturbed state ⟨*A*⟩_0_ to the final state:

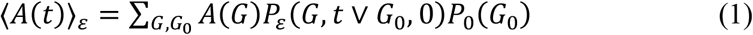

which can be interpreted as the average of all possible dynamical paths originating from initial configurations weighted by the probability distribution *P*_0_(*G*). The conditional probability to evolve from *G*_0_ to *G* is *P*_*ε*_(*G, t* ∨ *G*_0_, 0). Following the Onsager principle, the conditional probabilities are equal to those of spontaneous equilibrium in state 2. Hence, we can express them as the spontaneous fluctuations of state 1 (*P*_*ε*_(*G, t* ∨ *G*_0_, 0)) corrected by the presence of the perturbation term (*εB*(*G*)):

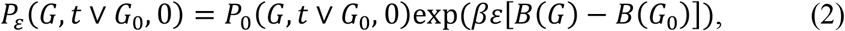

where *β* is the inverse temperature from equilibrium thermodynamics. Using this, the expectation value can be linearized and expressed as:

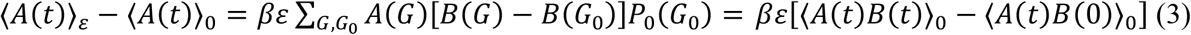

The susceptibility is defined as the variation of ⟨*A*⟩ in terms of the perturbation. We can define the time-dependent susceptibility as

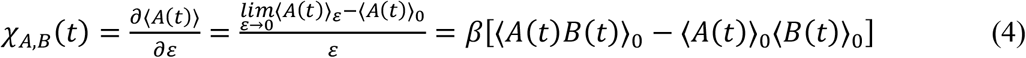

We can also obtain the static form of FDT by taking the limit *t* → ∞, obtaining the correspondence between the response of a system to a perturbation (on the left-hand side) and its unperturbed correlations (on the right-sand side) in equilibrium:

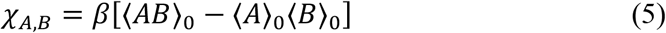

#### Violations of the FDT in non-equilibrium

If we describe the unperturbed state to have the mean values of the observables equal to zero (⟨*A*⟩_0_ ⟨*B*⟩_0_ = 0), then we can characterize the level of non-equilibrium as the normalized divergence of a system from the FDT:

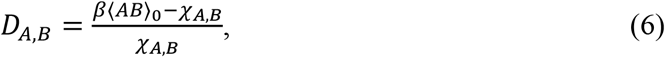

where *β* ⟨*AB*⟩_0_ explains the unperturbed fluctuations and *χ*_*A,B*_ corresponds to the response to a small perturbation *ε*. To obtain the total deviation of the FDT (*D*), we need to compute the mean value of *D*_*A,B*_ for all observables *A* and all perturbation points *B*. Thus, by leveraging the degree of deviation from the FDT we are assessing the distance from equilibrium of a system.

### Model-based FDT of whole-brain data

#### Whole-brain model

In order to estimate the total violation of the FDT (*D*, equation 6), we need to systematically perturb all brain regions *B* and observe the respective responses on all brain regions *A*. Thus, our initial step involves constructing a whole-brain model fitting the functional neuroimaging data in each group. This model allows us to derive the analytical expressions elucidating the correlations among all brain regions under spontaneous fluctuations and, subsequently, to apply a perturbation in a brain region and predict their average effect in all the other regions.

The whole-brain model consists of describing each node’s local dynamics as the normal form of a supercritical Hopf bifurcation, capable of defining the transition from asynchronous noisy behaviour to oscillations (84). Specifically, the whole-brain dynamics are represented by coupling the local dynamics of *N* = 80 Hopf-oscillators through the connectivity matrix *C*, which is given by:

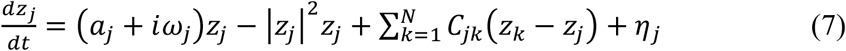

In this equation, *z*_*j*_ = *x*_*j*_ + *iy*_*j*_ represents a complex state variable of region *j* where *x*_*j*_ = *Real*(*Z*_*j*_)and *y*_*j*_ is the imaginary part. *η*_*j*_ is the additive uncorrelated Gaussian noise with variance *σ*^2^. The parameter *a*_*j*_ represents the node’s bifurcation parameter, with *a*_*j*_ < 0 describing the local dynamics as a stable fixed point at *z* = 0 and *a*_*j*_ > 0 creating a stable limit cycle oscillation with frequency *f*_*j*_ = *w*_*j*_/2 π. The intrinsic node frequency *w*_*j*_ is obtained from the empirical data by bandpass filtering the fMRI signals within the frequencies of 0.008–0.08 Hz and averaging the peak frequency of each brain region. We selected the parameter *a* = −0.02 based on Deco et al (84), as it generates a signal that is more effective at preserving resting state network structure and dynamically responding brain networks. The proximity to this critical point holds significant importance as it facilitates linearity in the dynamics. This linearity enables an analytical solution for the connectivity matrix *C*, which is determined by computing Pearson correlations between each pair of brain regions.

By employing a linear noise approximation (LNA) method, we can evaluate the functional correlations of the entire brain network. Consequently, the dynamical system involving N nodes (described by Equation 7) can be represented in a vectorized format:

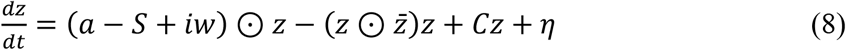

Here, *z* = [*z*_1_, …, *z*_*N*_]^*T*^, *a* = [*a*_1_, …, *a*_*N*_]^*T*^, *w* = [*w*_1_, …, *w*_*N*_]^*T*^, *η* = [*η*_1_, …, *η*_*N*_]^*T*^, and *S* = [*S*_1_, …, *S*_*N*_]^*T*^, which is a vector indicating the connectivity strength of each node, where *S*_*i*_ = ∑_*j*_ *C*_*ij*_. The transpose operation is denoted by []^*T*^, the ⊙ symbol represents the element-wise multiplication, and 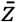 represents the complex conjugate of *z*. This equation describes the linear fluctuations around the fixed point *z* = 0, which is the solution of 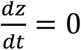. The progression of the linear fluctuations can be articulated using a Langevin stochastic linear equation by segregating the real and imaginary components of the state variables and disregarding higher-order terms 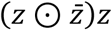:

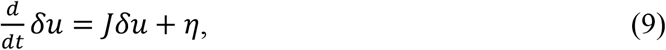

where the 2N-dimensional vector *δu* = [*δx, δy*]^T^ = [*δx*_1_, …, *δx*_*n*_, *δy*_1_, …, *δyn*]^T^ includes the fluctuations in both the real and imaginary aspects of the state variables. The 2*Nx*2*N*matrix *J* stands for the Jacobian of the system computed at the steady state and can be represented in the form of a block matrix:

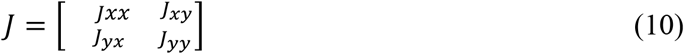

being *J*_*xx*_, *J*_*xy*_, *J*_*yx*_, *J*_*yy*_ matrices of size *NxN*. Specifically, *J*_*xx*_ = *J*_*yy*_ = *diag*(*a* − *S*) + *C* and *J*_*xy*_ = −*J*_*yx*_ = *diag*(*w*), where *diag*(*v*) represents a diagonal matrix with the vector *v* on its diagonal. Importantly, this linearization is only applicable if *z* = 0 is a stable solution, meaning that all eigenvalues of *J* have a negative real part.

#### Measuring violations of the model-based FDT

Using the described model, we can compute the violations of the FDT. First, we start by computing the covariance matrix *K* = ⟨*δuδu*^*T*^⟩. To do it, we first express Equation 9 as *dδu* = *Jδudt* + *dW*, where *dW* stands for a Weiner process with covariance ⟨*dWdW*^*T*^⟩ = *Qdt*. Here, *Q* is the covariance matrix of the noise. Then, using Itô’s stochastic calculus, we obtain *d*(*δuδu*^*T*^) = *d*(*δu*)*δu*^*T*^ + *δud*(*δu*^*T*^) + *d*(*δu*)*d*(*δu*^*T*^). Considering that ⟨*δudW*^*T*^⟩ = 0, maintaining the terms to first order in the differential *dt* and taking the expectations, we get:

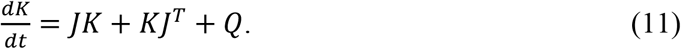

This allows us to compute the static covariance by

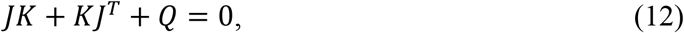

which can be solved using the eigen-decomposition of the Jacobian matrix *J* (85) and, from the first *N* rows and columns of the covariance *K* we finally get the functional connectivity simulated by the model (*FC*^*Simulated*^), which represents the real part of the dynamics.

In order to fit the model to the empirical data, we need to optimize the coupling matrix *C*, starting off the primer diffusion tensor imaging, such that the model accurately replicates the empirical covariances (*FC*^*empirical*^), computed as the normalized covariance matrix of the functional neuroimaging data, and the empirical time-shifted covariances (*FS*^*empirical*^(*τ*)), where *τ* denotes the time lag and are normalized by dividing each pair of nodes *i,j* by 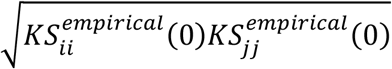. We select *τ* = 2, which is the value that minimizes the average autocorrelation. Interestingly, adjusting the timeshifted correlations can introduce asymmetries in the coupling matrix *C*, leading to non-equilibrium dynamics and violations of the FDT. The fitting is performed using a heuristic pseudo-gradient approach, iterating by updating *C* until it is completely optimized using:

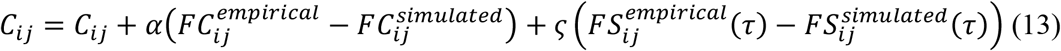

To compute *FS*^*Simulated*^(*τ*), we select the first N rows and columns of the simulated time-shifted covariances *KS*^*simulated*^(*τ*) normalized by dividing each pair of nodes *i,j* by 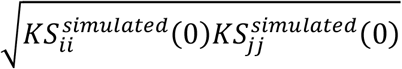. *KS*^*simulated*^(*τ*) represents the simulated time-shifted covariance matrix and is calculated as:

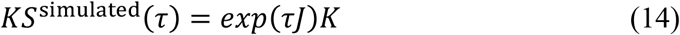

In the fitting of the model, convergence is reached when the updated value of *C* reaches a stable value, indicating that the model is optimized. *C* is initialized using the anatomical connectivity and only updated if there are connections in this matrix. This is followed strictly except in the homologous connections between the same regions in both hemispheres, which are also updated, as tractography is less accurate when incorporating this kind of connectivity. Moreover, *α* and *ς* are set to 0.00001. For each iteration, we calculate the model results by averaging the simulations corresponding to each participant. Finally, we refer to the optimized *C* as Effective Connectivity (EC) (34).

Having a fitted coupling matrix C for each participant and each group, we formulate an analytical expression of Equation 6 for the deviation from the FDT. Initially, we compute the expectation values of the state variables ⟨*δu*⟩_*εj*_when a perturbation *ε* is applied to a node *j*. Utilizing Equation 9, we can determine that 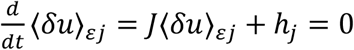, being *h*_*j*_ a 2*N* vector primarily of zeros except component *j* that equals the value of the perturbation *ε*. Solving for the desired expectation value, we obtain ⟨*δu*⟩_*εj*_ = −*J*^−1^*h*_*j*_. Considering only the real part of ⟨*δu*⟩_*εj*_, we obtain ⟨*δx*⟩_*j*_ = ⟨*δx* ⟩_*εj*_/*ε*. With this, we proceed to define the deviation from the FDT for region *i* when a perturbation is applied to region *j*:

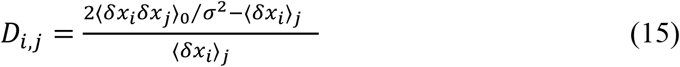

Here, the covariance ⟨*δx*_*i*_*δx*⟩_0_is computed from *KS*^*Simulated*^, and the term 2/*σ*^2^ is the inverse of temperature *β*. Then, the effect of perturbing region *j* on all the other regions can be computed as:

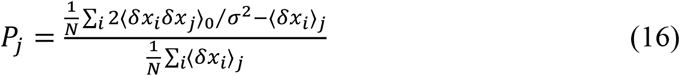

With this equation, we can define a vector, the perturbability map *P*, which represents the effect of perturbing over all brain regions. Using it, we can compute the level of non-equilibrium for each participant by averaging the deviation from the FDT over all possible perturbations:

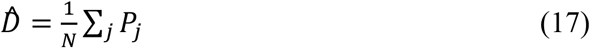

The analysis across the whole-brain network is performed both at the subject and at the node level. In the first case, we compute 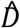 for each participant belonging to each group, obtaining one average value of FDT deviations of cortical nodes per subject. In the second case, we average the perturbability maps *P* across subjects in each group, obtaining one averaged *P* per group, which represents the mean value per node. Furthermore, to assess the variability of nodes, we compute the standard deviation of *P* of each subject in each group. To perform the RSNs analysis, we select the nodes belonging to each RSN and we do a subset of *P* only including them. Then, we perform the same analysis of the node level for each RSN: we average the subset *P* across subjects in each group, obtaining one averaged *P* per group. Moreover, we perform an analysis to understand the difference of the mean value of FDT deviation of each group in each RSN. To do this, for each RSN we compute the standardized mean difference of the averaged *P* of each group, that is, the subtraction of the means of the two groups divided by squared root of their standard deviations.

### Support Vector Machine Classification

Initially, we train an SVM classifier to distinguish between subjects in each group, using as input the FDT violations. Subsequently, we conduct an assessment of classifier performance using four distinct SVM linear models. These classifiers are trained on different input data, specifically: FC derived from empirical data, EC computed within the optimized whole-brain model, violations of the FDT, and a combination of EC and FDT violations. In both cases, the training process involves utilizing 80% of the dataset, while the accuracy is evaluated on the remaining 20% across 1000 k-folds. Furthermore, a detailed analysis is applied to the model exhibiting the highest accuracy to comprehend its feature importance. This involves extracting the coefficients of the model, sorting them from highest to lowest, and selecting the highest 10%. These top coefficients correspond to specific nodes within the brain parcellation, elucidating the significant brain regions driving the classification.

### Exploratory Factor Analysis

We follow the methodology in Schöttner et al (86) to reduce the dimensionality of our behavioral dataset. Using the “Factor Analyzer” package implemented in Python, we focus our analysis on variables from the Scale Analysis of Positive Symptoms (SAPS) and Scale Analysis of Negative Symptoms (SANS) that have less than 10% missing values. Prior to the analysis, all variables are standardized (z-scored) across subjects. The “Factor Analyzer” package facilitates dimensionality reduction by extracting underlying factors from the dataset. This process results in the identification of three factors, each comprising loadings for individual variables and scores for each subject. These factors represent distinct dimensions of symptomatology within our dataset.

### Statistical Analysis

To gain insight into the differences in FDT violations between individuals with schizophrenia and the control cohort, we employ the non-parametric independent-sample Mann-Whitney U test with 10000 permutations. To understand the relationship between FDT violations and symptom severity, we conduct correlation analysis between the average FDT violation value per subject and the scores obtained in each factor. In all analyses, significance is established for p-values below 0.05. For the analysis conducted between groups at the RSNs level, we implement the False Discovery Rate (FDR) correction method to account for multiple comparisons (87). The p-values outlined are the ones obtained after FDR correction. Preceding all the statistical analyses, outliers are excluded from the comparison, with outliers defined as data points deviating more than three standard deviations from the mean.

